# Integrative Multi-Omics Analysis of Melanoma: Uncovering Pathways Associated with Immunotherapy Outcomes

**DOI:** 10.1101/2025.04.01.646686

**Authors:** Morgan L. MacBeth, Alexander M. Kaizer, Brandie D. Wagner, Kasey L. Couts

## Abstract

**Purpose:** The treatment of advanced malignant melanoma with immune checkpoint blockade (ICB) therapies such as anti-CTLA4 and anti-PD1 has been transformative, yet a significant proportion of patients demonstrate intrinsic resistance or develop severe immune-related adverse events (irAEs), complicating treatment strategies. This study aimed to integrate clinical and molecular data using multi-omics factor analysis (MOFA) To better understand the multifaceted interactions governing ICB resistance and irAE development.

**Methods:** Melanoma patient-derived xenograft tumors with transcriptomic and microbiome data were analyzed using the MOFA2 R package. Simulations assessed MOFA2’s performance with small sample sizes. Transcriptomic and microbiome data were normalized and analyzed with MOFA2, and gene set enrichment analysis (GSEA) was performed.

**Results:** MOFA2 demonstrated robust performance with small sample sizes in simulations and accurately recapitulated findings from published data. Analysis identified five latent factors associating the tumor transcriptome, tumor microbiome, or both with differences in tumor subtypes, ICB response, and specific irAEs. GSEA highlighted pathways related to oxidative phosphorylation, DNA replication, and immune responses.

**Conclusion:** Integrative analysis of multi-omics data using MOFA2 provides insights into melanoma biology, uncovering distinct molecular pathways underlying clinical phenotypes. These insights contribute to our understanding of the complex biological mechanisms contributing to differences in melanoma clinical and tumor characteristics and treatment response, offering potential insight towards future development of more personalized and effective diagnostic, prognostic, and therapeutic strategies for patients.

**Context Summary:** This study aims to integrate multi-omics data, specifically transcriptomic and microbiome datasets, using the MOFA2 computational framework, to understand the complex interplay driving melanoma heterogeneity and response to immune checkpoint blockade (ICB) therapy. The study demonstrates that MOFA2 performs effectively even with small sample sizes, successfully capturing factors that distinguish tumor subtypes, ICB response, and immune-related adverse events (irAEs). It identifies associations between molecular features and clinical outcomes, shedding light on potential mechanisms underlying melanoma pathogenesis and treatment response. By integrating clinical and molecular data, the findings offer insights into the biological underpinnings of melanoma treatment response. Understanding these mechanisms could inform the development of more effective diagnostic, prognostic, and therapeutic strategies for melanoma patients, moving towards personalized oncology approaches.

## Introduction

Melanoma represents a formidable challenge in oncology due to its aggressive nature and proclivity for metastasis despite early detection. While immune checkpoint blockade (ICB) therapies including anti-CTLA4, anti-PD1, and their combination, have revolutionized the treatment of advanced malignant melanoma, nearly half of melanoma patients demonstrate intrinsic resistance to ICB treatment. [1–3] A subset of patients also develop severe immune-related adverse events (irAEs) in response to ICB, particularly combination anti-CTLA4 and anti-PD1, which can lead to delayed or discontinued treatment.

Numerous studies have analyzed clinical data and identified key features associated with ICB response and irAEs. For example, women are more likely to have a higher irAE burden with ICB treatment. [4] In addition to clinical data, the explosion of new and affordable high-throughput technologies such as transcriptomic, genomic, proteomic, and metabolomic assays, has provided researchers with vast amounts of molecular data. These conventional single-omics approaches have granted scientists an unprecedented look at the molecular machinery behind melanoma cellular functions including tumor immunity and ICB response. From these single-omics studies, several mechanisms have been identified which contribute to ICB resistance, such as loss of major histocompatibility complex (MHC) I/II expression, loss of phosphatase and tensin homolog (PTEN), and activation of the transforming growth factor beta (TGFβ) pathway. [5]

Despite vast amounts of clinical and molecular data on melanoma ICB response and irAEs, we have yet to develop an integrated landscape of the clinical and molecular factors driving ICB resistance and development of severe irAEs. Unfortunately, single-omics data often overlook the complex interplay between multiple molecular layers, limiting our ability to comprehensively understand the underlying mechanisms that govern physiological processes. Combining single-omics datasets, along with clinical information, could lead to novel associations and treatment strategies for disease. To date, few studies have performed unsupervised integration of clinical and molecular datasets for ICB-treated melanoma patients and tumors, yet these types of unbiased analyses could lead to new and unexpected insights into melanoma ICB response, related factors and irAEs. However, integrating these complex data presents unique challenges such as differing data dimensions and sample size, scale, accuracy, and conflicting modes. [6]

In recent years, the advent of multi-omics technologies to overcome the described challenges has revolutionized our ability to interrogate cancer biology at unprecedented depths by integrating genomic, transcriptomic, epigenomic, proteomic, and metabolomic data sets. Among these, multi-omics factor analysis (MOFA) has emerged as a powerful computational framework capable of unraveling the intricate relationships between different molecular layers and identifying key factors driving biological variability within and across cancer subtypes. MOFA seeks to uncover latent, or “hidden”, factors across datasets by reducing the dimensionality into groupings of variables to explain variation across samples. Combining these latent factors from - omics data with clinical features could be useful in explaining associations across samples not readily seen in individual -omics datasets.

A challenge of implementing MOFA is interpreting the resulting latent factors. MOFA2 is a recent R package that was proposed with modifications to attempt to address this challenge. Specifically, model inference is performed using variational Bayesian inference with mean-field assumption [7]. The resulting optimization problem consists of an objective function that maximizes the data likelihoods under some sparsity assumptions [8] which yields a more interpretable model output. In the remainder of the paper, when we are referring to analysis using the R package we will use ‘MOFA2’, and when referring to the general method we will use ‘MOFA’.

In this study, we utilize MOFA2 to integrate melanoma clinical patient and tumor features with multi-omics analysis of melanoma tumor transcriptomic and microbiome datasets. The melanoma patient tumors with both transcriptomic and microbiome data available represent variability in ICB response and patient features such as irAEs and demographics including gender, hair color, eye color, skin type, pre-existing co-morbidities. The data also include a subset of patient tumors of the rare mucosal melanoma subtype, which has been shown to have overall poor response and survival benefit with ICB compared to common cutaneous melanoma. [9–12] Given the smaller sample size of melanoma tumors with availability of clinical, transcriptomic, and microbiome data compared to previous datasets utilized for MOFA2 analysis, we first evaluate MOFA2 performance with a smaller sample size using previously evaluated data and simulation studies. Ultimately, through MOFA2-driven analyses, we seek to identify common and distinct molecular pathways, gene expression patterns, and microbiome alterations that may characterize cutaneous and mucosal melanoma, ICB response and/or irAE development. By shedding light on the underlying biological differences, we aim to advance our understanding of melanoma pathogenesis and anti-tumor immunity, and inform the development of more effective diagnostic, prognostic, and therapeutic strategies for patients afflicted with this deadly disease.

## Methods

### MOFA2

All analyses were performed using R Statistical Software (version 4.3.1, R Core Team 2023). The input to MOFA2 is a list of matrices with matching samples where each matrix represents a different data modality called a view and includes information called features. For example, RNA-sequencing and 16S ribosomal RNA (rRNA) sequencing would be defined as different modalities where genes and bacteria would be the features included, respectively. Data is normalized before running. After model fitting, the resulting factors are then characterized. Sample metadata such as demographics, mutational status, or treatment status is added at this stage to assist in characterization. For a second stage analysis, correlations between the factors and this sample metadata were estimated using the ‘correlate_factors_with_covariates()’ function provided in the MOFA2 package.

The formula for MOFA2 based on matrices is as follows:

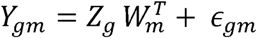

Where *Y* is the matrix of observations, *Z* is the factor matrix, *W* is the weight matrix, E is the residual error matrix, *g* indexes group, and *m* indexes modality. E can be specified to account for continuous, binary, or count data with Gaussian, Bernoulli, or Poisson models respectively. Model inference is executed using variational Bayesian inference with mean-field assumption. [8] Weights inferred from the model provide a score for how strongly each feature relates to each factor to allow a biological interpretation of the factors. A positive weight indicates the feature has higher levels in the samples with positive factor values and negative weights indicate the feature has lower levels in the samples with negative factor values. Features that have no association with factors have weights closer to zero, and features with strong association with the factor will have large absolute weight values.

### Datasets used to evaluate small sample size performance using MOFA

Data were simulated using a modified built-in data generation function from the MOFA2 R package. A large model was specified that included 1000 samples, 2 views with 2500 and 150 features representing the ‘true’ model used as a benchmark to evaluate performance in smaller subsamples. The sampled models had 25 samples each to mirror our real-world melanoma data. Models were compared by taking the top 50 features in each view by weight assigned in the model and comparing overlapping features in each factor across both views in the ‘true’ model and the sampled models. Factor labels could switch in the smaller subsamples, so alluvial plots were produced to aid in the visualization of the top 50 features regardless of the factor label. The weights of the features in each factor were used to rank the features and relate them to the features and factors present in the original model.

In addition, 28 samples were randomly selected from a published study using MOFA including viral, fungal and bacterial sequence data to evaluate performance in smaller sample sizes. [13] Metadata from the study was used to generate graphs and compare factors found using the full set of samples in the original study.

### Application of MOFA to melanoma transcriptome and microbiome data

Melanoma transcriptome TPM data was obtained from total RNA sequencing of patient-derived xenograft models available in dbGap under accession number phs002951.v1.p1. Bacterial 16S rRNA data was generated directly from melanoma patient tumors as previously described (manuscript under review).

Bacterial 16S rRNA ASVs and RNA sequencing TPMs were defined as different data modalities. 16S sequencing counts and RNA sequencing transcripts per million (TPM) were normalized using a centered log ratio (CLR) transformation before a MOFA model was generated. [14] CLR has been shown to be effective in normalizing compositional data. [15] 16S data was used at the order level and filtered to exclude features with 0 counts across all samples prior to CLR transformation. RNA sequencing data was restricted to the top 2500 features across the 28 samples and excluded pseudogenes. The number of factors was changed from the default of 10 to 5 in the MOFA model options and ‘slow’ convergence mode was used.

### GSEA

Gene set enrichment analysis was performed in the MOFA2 package using the Molecular Signatures Database C2 (MSigDB) gene set annotation. Separate analyses were performed for genes with positive and negative weights. A parametric t-test was the statistical test used to compute significance of the gene set statistics under a competitive null hypothesis.

## Results

### MOFA2 identifies similar ground truths with smaller sample sizes

We first sought to test the MOFA2 R package performance with a small number of samples using simulated data. The built-in simulation function in the MOFA2 R package did not allow for a differing number of features in each view, therefore it was modified to simulate data that better reflects our melanoma dataset and is included in the supplementary data. Using this modified function, data was simulated for 1000 samples, 2 views, and 5 factors (**Fig. 1A**). One view had 2500 features and one had 150 features, similar to our melanoma datasets. A MOFA2 model using this data was used as the ‘true’ model with which to compare factors estimated from smaller subsets. A subset of 25 samples was randomly sampled from the ‘true’ data and again a MOFA2 model was fit (**Fig. 1B**), and this was repeated 1000 times. The Jaccard similarity index score between top features, as determined by weight, across the five factors was calculated as a measure of agreement. Percent agreement between features in each factor was also reported (**Fig. 1C,D**). Greater than half of all features in each factor were overlapping according to the Jaccard similarity score (**Fig. 1C**). The percentage agreement of features shared among the five factors in the ‘true’ and sampled models across the 1000 simulations showed between 58%-74% agreement, with the greatest similarity in Factor 1 and least similarity in Factor 5 (**Fig. 1D**). It is worth noting that features are not unique to factors and can be shared among them, which is why percentages in Figure 1D do not add up to 100. Finally, two representative simulations were randomly chosen from the 1000 sampled and alluvial plots were created for each to evaluate how features in the original factors from the ‘true’ data compared to the smaller sampled models. The alluvial plots revealed that many of the top features are shared among the same factors in the ‘true’ model and the smaller sampled model, with only a small subset of features moving to different factors in the smaller model (**Fig. 1E**). Overall, MOFA2 was able to recapitulate the majority of features in all factors from the ‘true’ model across the 1000 smaller sampled models.

**Figure 1.**
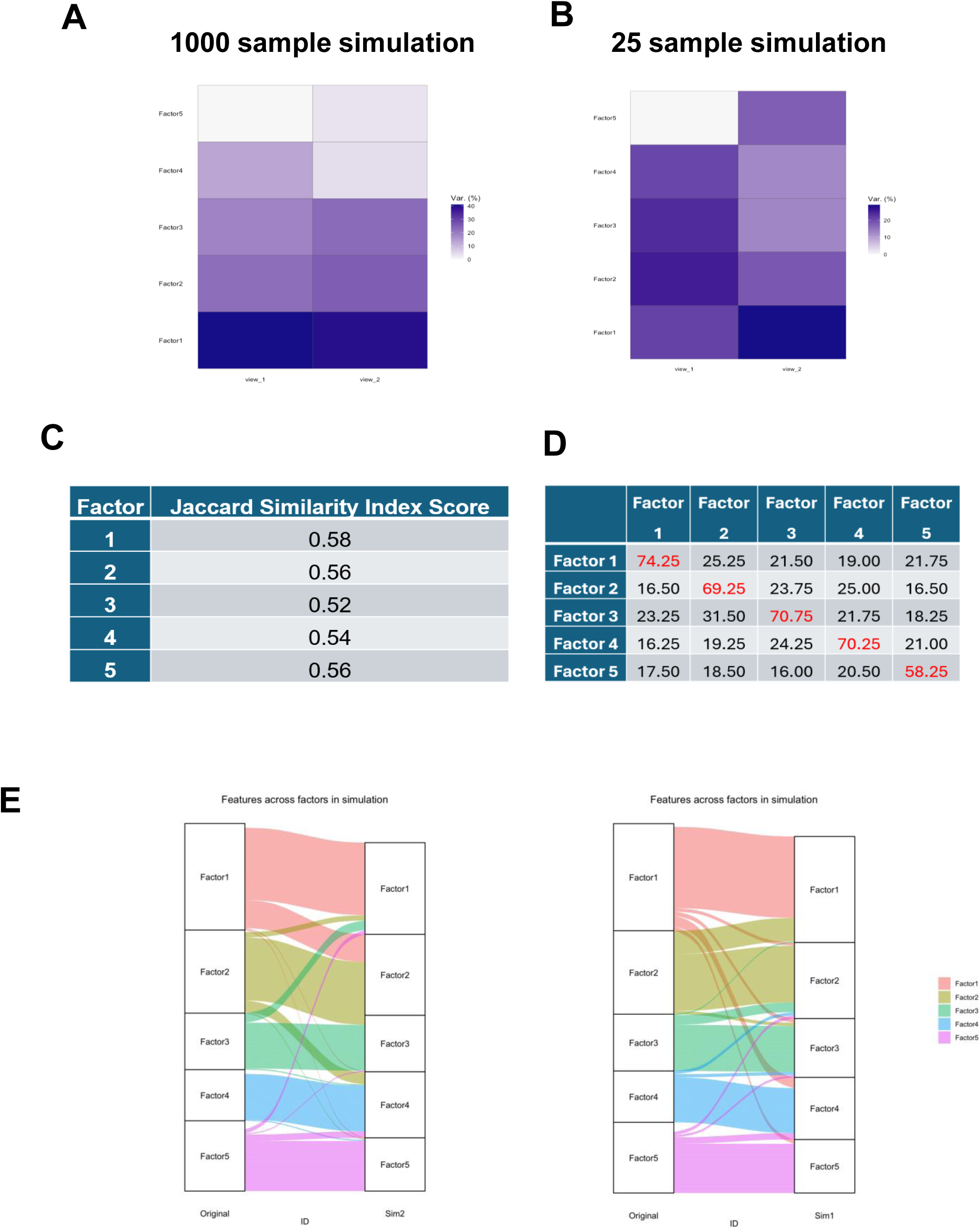
MOFA2 is able to recover ground truth from simulated data. A. Variance decomposition heatmaps for a representative sample and B. simulated dataset. C. Jaccard Similarity Index score and D. percent agreement table. Features can appear in more than one factor explaining why the rows and columns do not sum to 100 in the percent agreement table. E. Alluvial plots illustrating feature drift among factors for two representative samples. The left column is the ground truth from the simulated data and the right column is the sampled data.

### MOFA2 recaptures findings from published study with smaller sample

We next acquired data from a published study that was previously used as an example application of the MOFA2 package to further evaluate its performance in a real-world setting upon sample size reduction [13]. The original study included 59 samples with 16S rRNA, Intergenic Transcribed Spacer (ITS), and virus discovery next generation sequencing as well as metadata for these samples. The samples included stool samples from: 33 patients admitted to ICU and treated with antibiotics; and 26 healthy controls where 6 were treated with antibiotics and 20 were not treated. We randomly selected 28 samples to mimic the sample size of our melanoma dataset (**Fig. 2A**), and a MOFA model was fit and compared to the original findings. The original model identified 6 factors, while the model fit to the smaller subset identified 5 factors (**Fig. 2B**). In the original model, Factor 1 and Factor 2 had the most variance explained in the bacteria and fungi views, respectively, we found that also to be true in the smaller model. Factor 1 was strongly weighted by short chain fatty acid differences as measured by fecal metabolites butyrate, acetate, and propionate with log 10 adjusted p-values up to 8 in the original study. This was also recapitulated with the smaller sample cohort with p-values around 3 (**Fig. 2C**). Further, the original study also found microbial differences between study participants that received antibiotics and those who did not, reflected by plotting Factor 1 vs Factor 3 (**Fig. 2D**, left panel). In our reduced sample size dataset, MOFA2 also demonstrated similar sample clustering upon Factor 1 vs Factor 3 comparison (**Fig. 2D**, right panel). Altogether, MOFA2 was able to recapture factors found in published data as well as ground truth from simulated data in smaller sample sizes, providing confidence that MOFA could be applied to our similarly sized melanoma dataset.

**Figure 2.**
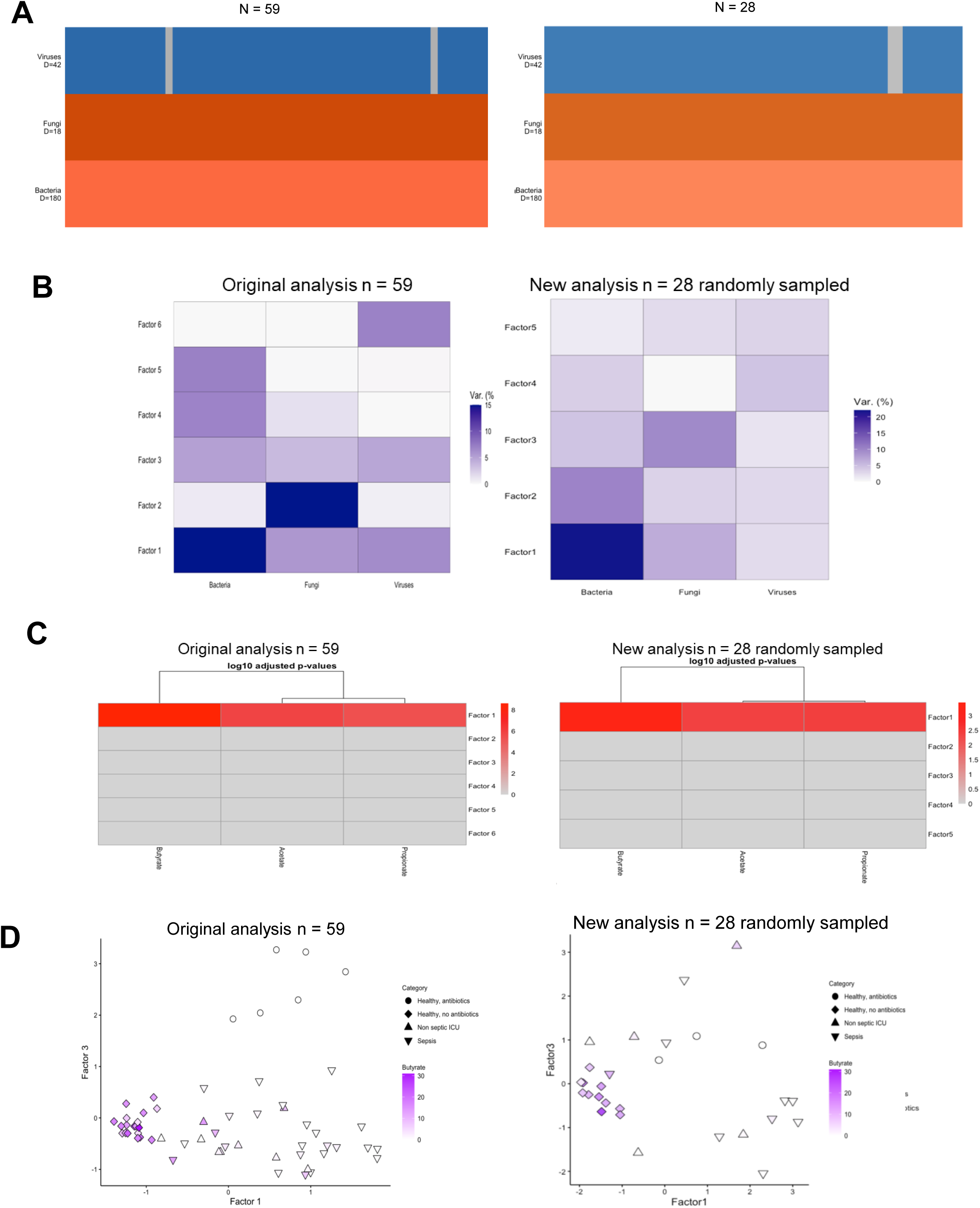
MOFA performance in a real-world dataset upon sample size reduction. Comparison of published dataset to sampled data looking at variance decomposition (A.), p-value heatmaps for correlation of fecal metabolites with factor 1 (B.) breakdown of samples with gray denoting a missing matched sample (C.), and scatterplot of factor weights in factor 1 vs factor 3 where point shape denotes sample category and color denotes butyrate concentration (D.).

### Clinical and genomic features of melanoma tumors included in analyses

There was a subset of 28 melanoma samples, 16 cutaneous and 12 mucosal melanoma tumor subtypes, which had both RNA sequencing and 16S rRNA data available (**Table 1**). Mean ages were similar for both subtypes, with a mean age of 61 (13.2 standard deviation (SD)) for cutaneous and 63 (14.2 SD) for mucosal. Of the 16 cutaneous melanoma samples, 9 were from nodular tumors, 6 from superficial spreading tumors, and 1 from melanoma in-situ. Among the corresponding cutaneous melanoma patients, 11 had received ICB with 5 responding to treatment and 5 non-responders and 1 response unknown, with the remaining 5 patients not receiving any ICB treatment. Among the 12 mucosal melanomas, 5 of the samples were from anorectal tumors, 3 from sinonasal tumors, and 4 from vulvovaginal tumors. All 12 of the corresponding mucosal patients had received IBC treatment, with 8 patients not responding and the remaining 4 did not have response data available.

**Table 1.** Clinical characteristics of melanoma samples.

|  | **Cutaneous**<br>**(N=16)** | **Mucosal**<br>**(N=12)** | **Overall**<br>**(N=28)** |
| --- | --- | --- | --- |
| **ICB Treatment Status** |  |  |  |
| On | 3 (18.8%) | 2 (16.7%) | 5 (17.9%) |
| Post | 3 (18.8%) | 5 (41.7%) | 8 (28.6%) |
| Pre | 7 (43.8%) | 4 (33.3%) | 11 (39.3%) |
| Missing | 3 (18.8%) | 1 (8.3%) | 4 (14.3%) |
| **Subtype** |  |  |  |
| Cutaneous - MIS | 1 (6.3%) | 0 (0%) | 1 (3.6%) |
| Cutaneous - Nodular | 9 (56.3%) | 0 (0%) | 9 (32.1%) |
| Cutaneous - SS | 6 (37.5%) | 0 (0%) | 6 (21.4%) |
| Mucosal - Anorectal | 0 (0%) | 5 (41.7%) | 5 (17.9%) |
| Mucosal - Sinonasal | 0 (0%) | 3 (25.0%) | 3 (10.7%) |
| Mucosal - Vulvovaginal | 0 (0%) | 4 (33.3%) | 4 (14.3%) |
| **Age** |  |  |  |
| Mean (SD) | 61.0 (13.2) | 63.6 (14.2) | 62.1 (13.4) |
| Median [Min, Max] | 59.0 [32.0, 84.0] | 64.5 [41.0, 87.0] | 61.0 [32.0, 87.0] |
| **ICB Response** |  |  |  |
| NR | 5 (31.3%) | 8 (66.7%) | 13 (46.4%) |
| R | 5 (31.3%) | 0 (0%) | 5 (17.9%) |
| Missing | 6 (37.5%) | 4 (33.3%) | 10 (35.7%) |
| **Received ICB** |  |  |  |
| N | 5 (31.3%) | 0 (0%) | 5 (17.9%) |
| Y | 11 (68.8%) | 12 (100%) | 23 (82.1%) |

### MOFA2 analysis identifies 5 latent factors associated with underlying various clinical metadata

An exploratory analysis using MOFA2 was performed to identify novel associations between the tumor microbiome and transcriptome which relate to clinical metadata. MOFA2 identified 5 latent factors, with the mRNA view explaining more variance in every factor than the bacteria view, which was predominant in Factors 1 and 4 (**Fig. 3A**). Correlation plots, summarized in **Fig. 3B**, were generated to identify clinical and tumor features associated with the latent factors. Metadata included demographics, comorbidities, tumor characteristics, response to immunotherapy, and immune-related adverse events (irAEs). Overall, factors were found to be significantly associated with features of tumor characteristics (**Fig. 3C**), immunotherapy response (**Fig. 3D**), and/or irAEs (**Fig. 3E**), with virtually no associations with patient-focused metadata of demographics or co-morbidities (**Supplementary Fig. S1**).

**Figure 3.**
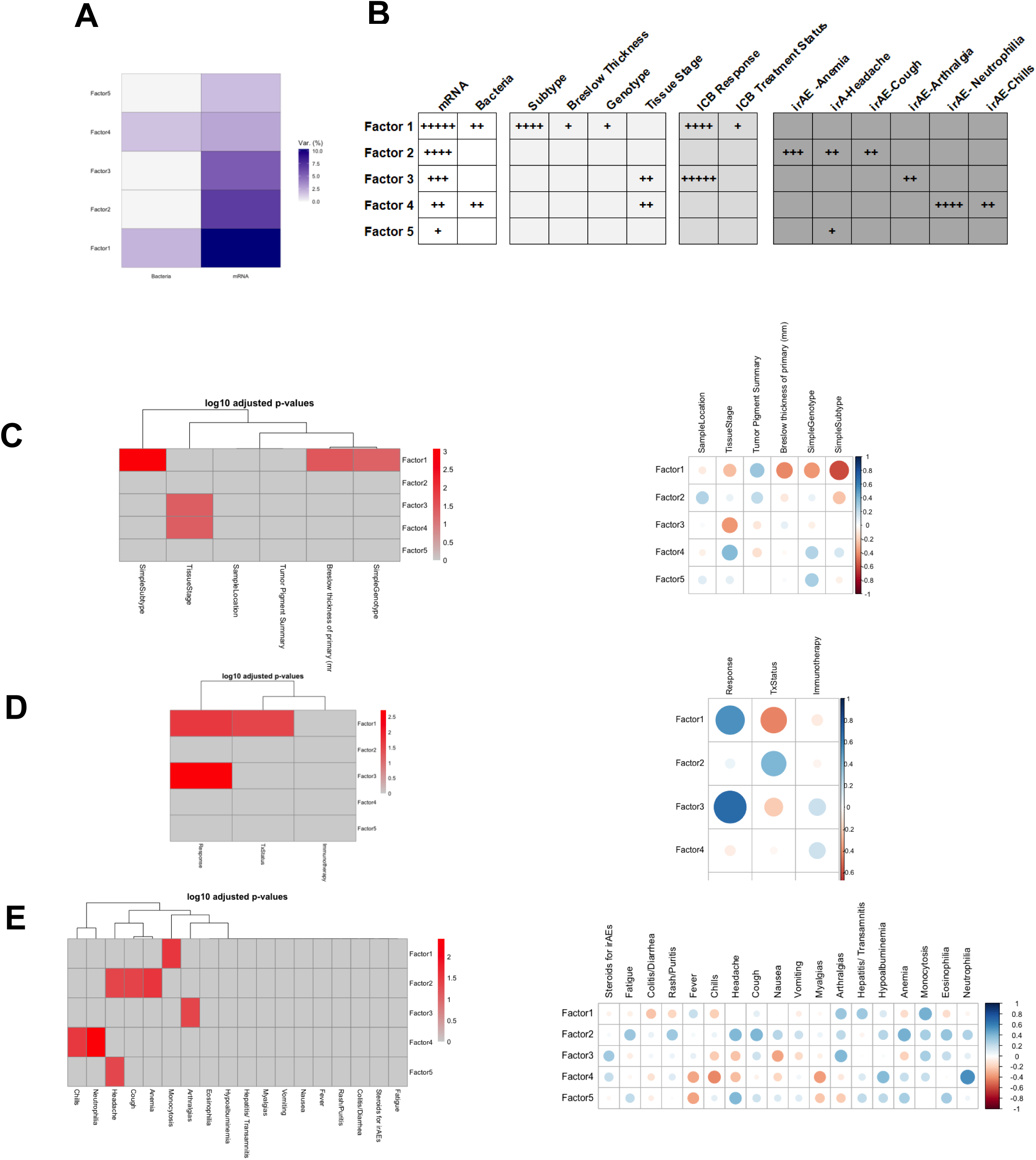
MOFA2 factors relate to distinct tumor and immune features. A. Variance decomposition heatmap across all five factors and two views. The darker the cell, the more variance explained by the model. B. Summary of variance explained on the far left with summary of significantly correlated tumor and immune features to the right. ‘+++++’ being the most highly correlated, and ‘+’ less but still significantly correlated. C. Correlation plots and p-value heatmaps of tumor characteristics, response (D.), and irAEs (E.). The darker the red in the heatmap, the more significantly correlated. Bold black boxes denote the most correlated variable on the plot.

Factor 1, including both mRNA and bacteria views, was significantly correlated with tumor features of subtype, Breslow thickness, and genotype (**Fig. 3C**), as well as immunotherapy response (**Fig. 3D**), and the irAE monocytosis. Factor 2, mRNA focused, was correlated with irAEs anemia, headache, and cough (**Fig. 3E**). Factor 3, mRNA focused, was significantly correlated with response to immunotherapy (**Fig. 3C**), with lesser correlation with tumor characteristic tissue stage (**Fig. 3C**) and irAE arthralgia (**Fig. 3E**). Factor 4, which included both mRNA and bacteria views, was significantly correlated with irAEs neutrophilia and chills, and weakly correlated with hypertension (**Fig. 3E, Supplementary Fig S2)** and tumor characteristic tissue stage (**Fig. 3B**). Finally, factor 5 was significantly correlated with the irAE headache (**Fig. 2E**). Overall, we saw mRNA explaining variance in every factor, and bacteria only explaining a small amount of variance in two factors. Immunotherapy response significantly correlated with two factors and at least one of seven different irAEs significantly correlated with each of the factors.

### Distinct gene and/or bacteria features define each Factor

The MOFA2 model assigns weights to each feature in each latent factor. These weights correspond to how strongly a feature relates to a factor and provide a biological interpretation of the factors. Therefore, to identify the specific gene and/or bacteria molecular features contributing to individual factors, heatmaps showing gene expression values or bacterial counts for the genes or bacteria with the highest weights were generated for the mRNA and bacteria views to show distinct subsets that were either up- or down-regulated between clinical covariates. These specific gene set differences by factor will be explored in more detail in later sections. Factor 1, driven by both mRNA and bacteria features, most strongly separated melanoma subtypes and ICB responders (**Fig. 3C,D**), with the top 20 genes (13 higher in mucosal and 7 lower in mucosal) shown in **Fig. 4A**. Factor 1 subtype variance was explained with the top 20 bacterial classes shown (**Fig. 4B**). Mucosal melanoma samples had higher expression of *Firmicutes Clostridia*, an anaerobic bacteria known to be more common to gut and mucosal sites than skin, compared to cutaneous melanoma, whereas cutaneous melanomas included more *Actinobacteria* classes of bacteria. Factor 1 had also separated ICB responders in addition to melanoma subtype (**Fig. 3D**). Mucosal melanomas had higher expression of orders *Bifidobacteriales* and *Burkholderiales*, Gram-negative bacteria also known to colonize the gastrointestinal tract. In Factor 1, higher levels of anaerobic bacteria order *Clostridia* were correlated with ICB responses of non-responders (**Fig. 4B**). Factor 1 explained the most variance in the data and incorporated both mRNA and bacteria views. This factor revealed distinct mRNA and bacteria differences in melanoma subtype as well as ICB response.

**Figure 4.**
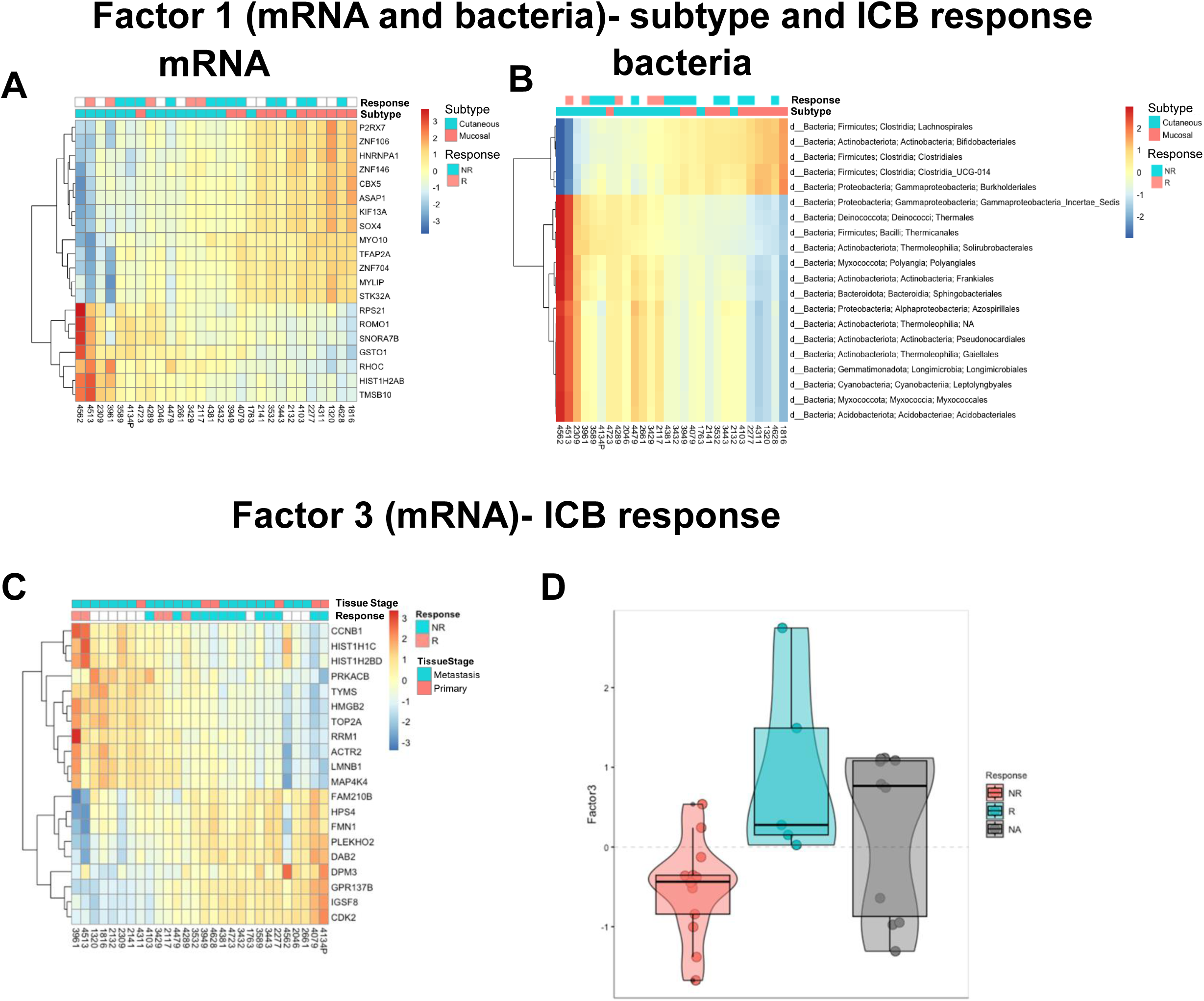
Molecular features defining Factors 1 and 3 in subtype and ICB response. A. Heatmap of top mRNA and B. bacteria features as selected by weight in factor 1 with corresponding tumor subtype and ICB response shown above. Red values correspond to more positive factor weights and blue values correspond to negative factor weights. C. Heatmap of top mRNA features as selected by weight in factor 3 with corresponding ICB response and tissue stage shown above. D. Violin plot of factor 3 weights on the y-axis by corresponding ICB status.

Factor 3 also separated immunotherapy responders and non-responders, but only based on mRNA, and was more highly correlated than Factor 1 with response. Unlike Factor 1, Factor 3 was specific to immunotherapy response and did not correlate with subtype, representing potential cutaneous melanoma specific features determining ICB response. The top 20 genes by weight with sample response information included above the heatmap are shown in **Fig. 4C**. Interestingly, the two distinct groups of immunotherapy responders and non-responders are shown clearly when plotting just the weights associated with the features in Factor 3 (**Fig. 4D**). For samples without this response information, there are two distinct groupings within this comparison suggesting a mixture that may correspond to responding (samples below the dotted line) or non-responding (samples above the dotted line) were data available (**Fig. 4D**, “N/A” group).

Whereas Factors 1 and 3 were associated with melanoma subtype and ICB response, Factors 2, 4, and 5 were mostly associated with specific irAEs. Factors 2 and 5 contained mostly mRNA features, and Factor 4 was made up of both mRNA and bacteria (**Fig. 3A,B**). Factor 2 was associated with anemia (a condition that develops when your blood produces a lower-than-normal amount of healthy red blood cells), headache, and cough, with the top 20 genes shown in **Fig. 5A**. Associated with neutrophilia (a higher-than-normal number of neutrophils) and chills the top 20 genes for Factor 4 are shown in **Fig. 5B**. In contrast to Factor 2, where features were consistent between all irAEs, Factor 4 associated features, neutrophilia and chills, were nearly mutually exclusive. Factor 4 bacteria features (top 20 shown in **Fig. 5C**) which were also nearly mutually exclusive between the two irAEs, included higher levels of the bacteria orders *Burkholderiales*, *Corynebacteriales*, and *Lactobacillales* in samples associated with neutrophilia, and higher *Deferribacterales*, *Obscuribacterales*, *Acetobacteriales*, and *Propionibacteriales* bacteria orders in samples associated with chills (**Fig. 5C**). These same bacteria and genes were lower in those with hypertension and chills as well as higher in those with neutrophilia. (**Supplementary Fig. S2B**) Factor 5 explained the least variance, but slightly separated those with headache. (**Supplementary Fig. S3A**)

**Figure 5.**
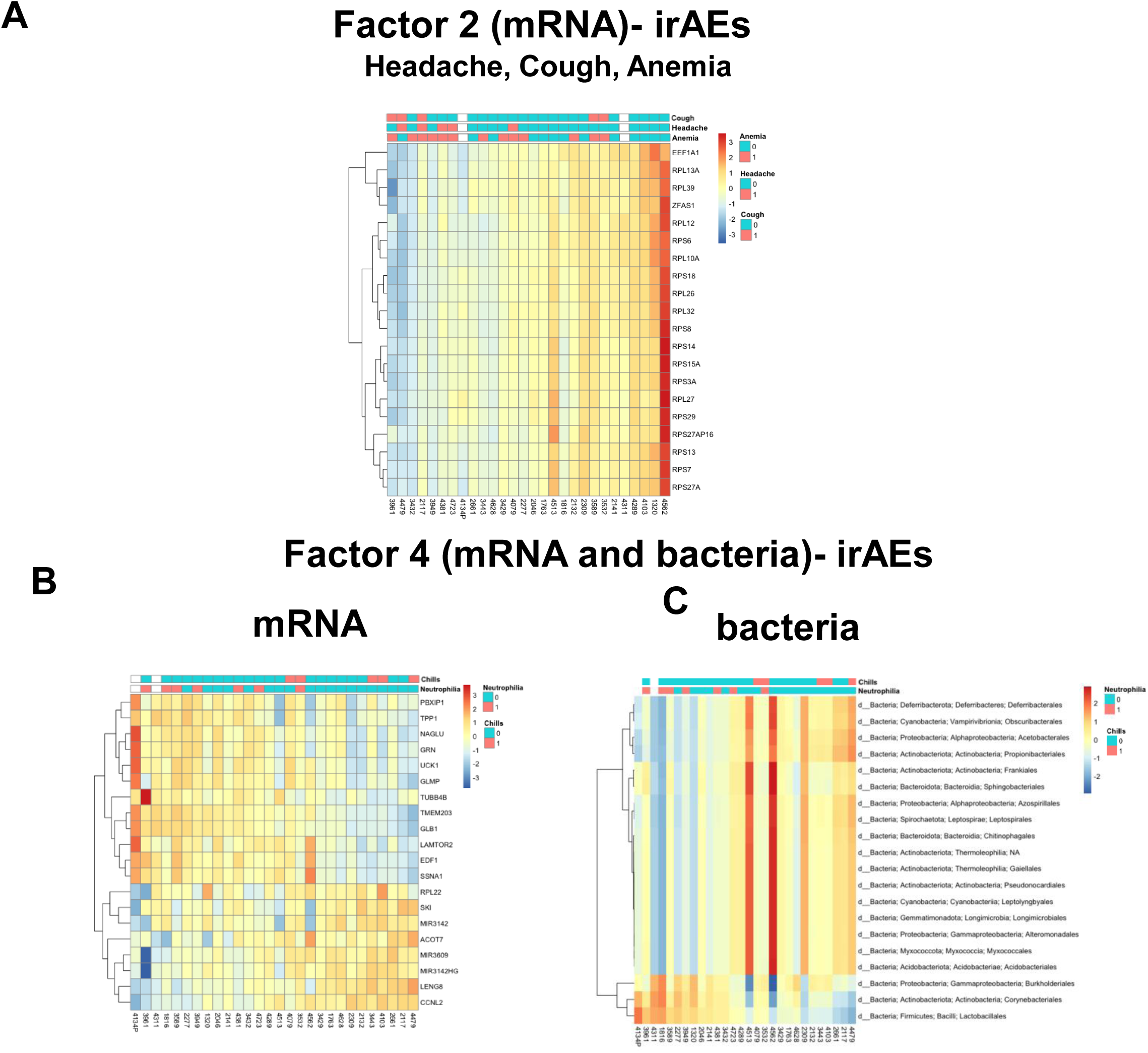
A. Heatmap of top mRNA features as selected by weight in factor 2 with corresponding irAE status above. B. Heatmaps of top mRNA and bacteria features as selected by weight in factor 4. Corresponding irAE status shown above heatmaps.

### *GSEA pathway* analysis to infer biological significance of Factor mRNA features

In order to investigate the MOFA factors in the mRNA view in more detail, gene set enrichment analysis (GSEA) was undertaken using the weights assigned in the MOFA model and the Molecular Signatures Database C2 (MSigDB) (**Fig. 6**). [16, 17] The C2 curated gene sets are from online pathway databases, knowledge of domain experts, and publications in PubMed. MOFA has positive and negative weights, with positively weighted features being ‘high’ in the samples with positive factor values, and negatively weighted features being ‘high’ in the samples with negative factor values. Therefore, separate analyses were performed for the positive and negative weights. A heatmap showing the gene sets included in the MSigDB C2 in rows versus factors in columns where each entry corresponds to the log p-value is shown in **Fig. 6A**. Heatmap results were converted into pathway enrichment plots for each Factor (**Fig. 6B, C**). For the positively weighted mRNA features in Factor 1, which defined both melanoma subtype and ICB response, the top pathways included oxidative phosphorylation and respiratory electron transport chain pathways. Factor 3 genes, defining only ICB response, were enriched for pathways of DNA replication, mRNA splicing, and cell cycle. In Factor 2, defining irAEs, the top pathways included the unfolded protein response and activation of chaperone genes by XBP1, which both deal with endoplasmic reticulum (ER) stress. Factor 4 mRNAs, differentially associated with neutrophilia and chills, showed enrichment of oxidative phosphorylation respiratory electron transport pathways which were interestingly similar to Factor 1. Factor 5, describing the least variance and only weakly associated with one irAE, had pathways associated with metabolism and translation (**Supplementary Fig. S3B**). For the negatively weighted features in factor 1, the peroxisome proliferator-activated receptor alpha (PPARA) pathway and inositol phosphate metabolism pathways were the top by negative log p-value. Top negatively weighted feature pathways for Factor 2 included peptide chain elongation pathways, mRNA metabolism, and nonsense mediated decay pathways. Iron uptake transport and lysosome were the top pathways for negatively weighted features in factor 3. Top pathways in the factor 4 negatively weighted features were transport of mature transcript to the cytoplasm, toll-like receptor 4 (TLR4) signaling, SMAD2/3:SMAD4 transcriptional activity, and the interferon gamma (IFNg) pathway. Finally, in Factor 5, the top pathways for the negatively weighted features mostly all dealt with RNA Polymerase II transcription. (**Supplementary Fig. S3C**) Interestingly, in the Factors 1 and 4 incorporating both the mRNA and bacteria data, there was an overlap of pathways such as oxidative phosphorylation, electron transport, Huntington’s disease, and Alzheimer’s disease. These two factors explained both ICB response and irAEs, respectively, suggesting a connection between the transcriptome and microbiome.

**Figure 6.**
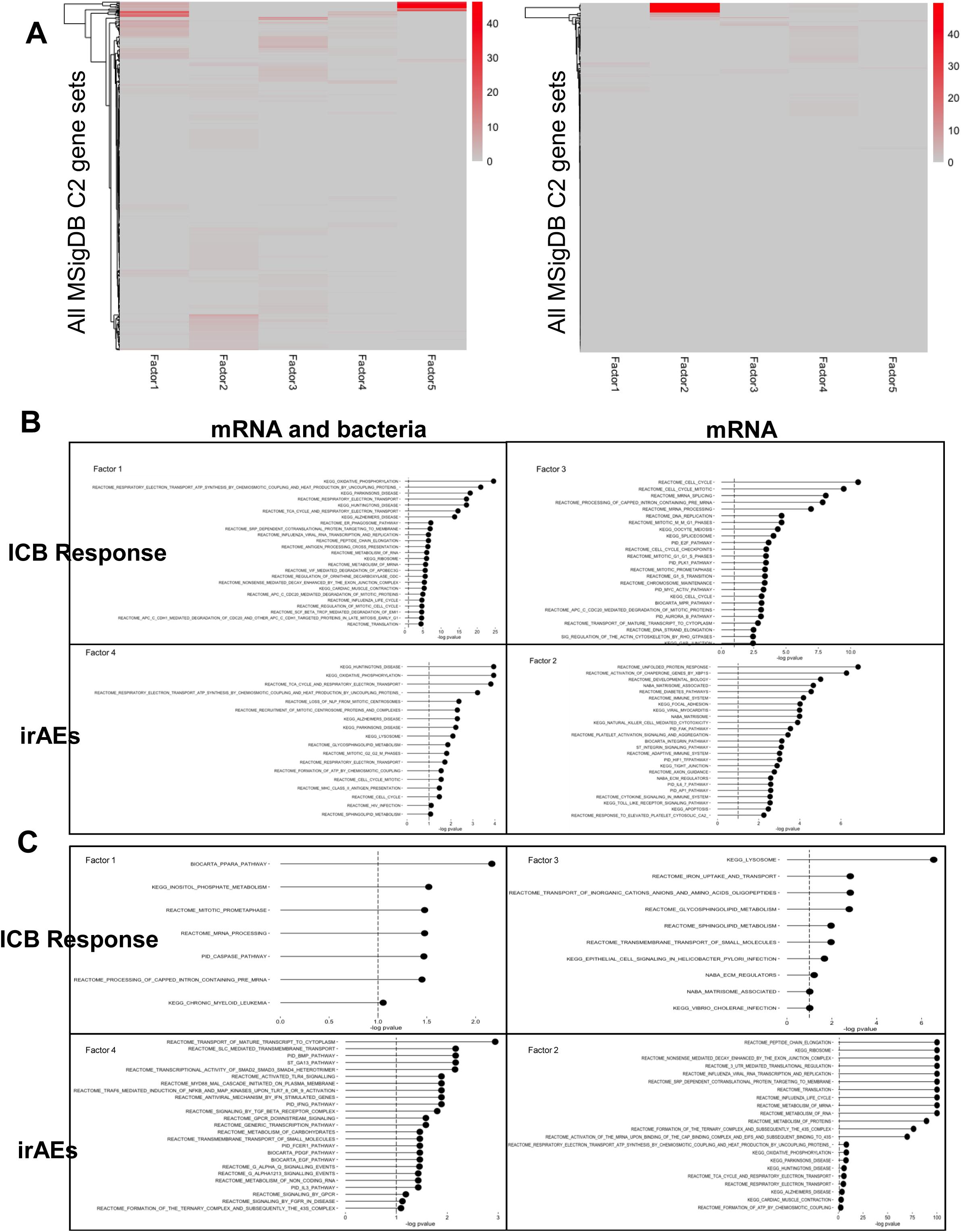
GSEA reveals unique gene expression pathways for each factor. A. A heatmap showing all gene sets included in the MSigDB C2 database (rows) versus factors (columns) where each entry corresponds to the log p-value. Statistically significant pathways are highlighted in red. Positive factor loadings shown on the left and negative factor loadings shown on the right. B. Top significant pathways for the positive feature weights in factors 1-4. C. Top significant pathways for the negative feature weights in factors 1-4.

## Discussion

The integration of multi-omics data has emerged as a powerful approach in cancer research, offering insights into the complex interplay between various molecular factors underlying clinical and tumor differences. [6, 18] In this study, we employed Multi-Omics Factor Analysis (MOFA) to synergistically analyze melanoma tumor microbiome and transcriptomic data features contributing to clinical and tumor characteristics. We selected MOFA since other approaches such as Multiple co-inertia analysis or Regularized Generalized Canonical Correlation Analysis may be limited because they consider the factors to be different for each -omics layer. In a recent review, MOFA was the only multi-omics approach that could detect slightly nonlinear signals. [18–20]. Our findings shed light on the intricate relationships between the tumor microbiome and transcriptome, and how they correlate with clinical features and treatment responses.

Our integrative analysis revealed novel associations between microbial composition and gene expression patterns, unveiling potential microbial drivers of tumor heterogeneity in cutaneous and mucosal melanoma. Through the identification of co-varying omics factors, we delineated intricate relationships between microbial taxa, host pathways, and clinical phenotypes, providing mechanistic insights into tumor-microbiome interactions.

Despite the increasing availability of multi-omics studies, these datasets can have small sample sizes. Previous simulation studies evaluating MOFA assumed relatively large sample sizes and a consistent number of features in each data modality being assessed, which may not be a reasonable assumption depending on the -omics platform. Before applying MOFA2 to our data, we evaluated its performance under conditions of a smaller sample size and differing number of features. The simulations range from ideal to close to real-world conditions based our melanoma data sample size as a motivating example. MOFA2 was able to recapture factors found in published data as well as ground truth from simulated data, leading us to have confidence in applying MOFA to our data.

After applying MOFA to our motivating dataset, factors 1 and 4 were potentially the most interesting as they had variability explained by both views in the MOFA model, while factors 2, 3, and 5 only incorporated the mRNA view. Factor 1 was more of a confirmatory factor since it was able to distinguish subtype and response differences. Factor 1 also explained the most variance of the factors, showing that subtype and response differences are driven by both mRNA and bacteria differences. With our small sample set we did not have any mucosal melanoma samples that were responders, so this could be a confounding variable and should be investigated bearing that in mind. Factors 3 and 4 correlated with tumor tissue stage, but only a small number were many primary tumor samples, so it is difficult to make conclusions about this correlation. No factor was correlated with demographic data and only one comorbidity had correlation with any factor, which could possibly be explained by the fact that these were tumor samples and not germline samples. As seen with the simulation study, the higher order factors had more variability with small sample sizes, so conclusions gained from these factors may need more validation in the future.

Even with a small sample size, MOFA2 was able to distinguish distinct known differences such as subtype between samples in factor 1. This factor was also correlated with Breslow thickness and tumor genotype, which are reflective of the cutaneous or mucosal melanoma subtype. [9, 12, 21] This ‘confirmatory’ factor served as a valuable positive control and led us to have confidence in results when examining the remaining factors.

While only incorporating the mRNA view, factors 2 and 3 were both correlated with irAEs and factor 3 was correlated with ICB response as well as tumor tissue stage. Factor 2 was correlated with irAEs headache, anemia, and cough suggesting these adverse events are related have similar gene expression. Pathways that were found in the positive factor loadings in factor 2 were associated with the immune system, protein modification, and hypoxia inducible factor (HIF) transcription factor pathway which recapitulates what has been found to be associated with anemia. [22] Factor 3 was correlated with the irAE arthalgia and this irAE had the same gene expression trends as responders to ICB treatment. It has been observed that metastatic melanoma patients that develop an irAE after ICB treatment usually have better overall survival, suggesting that in our data developing arthralgia could be a sign of a better response. [23]

Factor 4 was significantly correlated with the irAEs neutrophilia and chills in both the mRNA and bacteria views. Interestingly, neutrophilia was positively correlated while chills was negatively correlated with this factor. Neutrophils use oxygen-dependent processes to kill microorganisms, leading to the production of reactive oxygen species (ROS). Neutrophils make energy through glycolysis canonically, and top pathway found by GSEA in the positive feature weights of factor 4 was oxidative phosphorylation, suggesting perhaps a dysregulation in these samples. [24, 25]

Overall, our findings underscore the importance of integrative multi-omics approaches in deciphering the molecular basis underlying clinical and tumor differences and uncovering new avenues for precision oncology. By illuminating the crosstalk between the microbiome and transcriptome with this exploratory analysis, we move closer to unraveling the intricate landscape of melanoma tumor-host interactions which may better guide future development of personalized therapeutic strategies tailored to individual patients.

## Acknowledgements

We thank the patients and their families for sharing tissue and clinical information with the International Melanoma Biorepository and Research Laboratory.

---

**Supplementary Figure S1.** Tumor-data derived MOFA Factors not defining for patient co-morbidities. Correlation plots for demographic data (A) and comorbidities (B) across the factors.

**Supplementary Figure S2.** Additional Factor 4 mRNA and bacteria heatmaps. Heatmaps of top mRNA and bacteria features as selected by weight in factor 4. Corresponding hypertension status shown above.

## References

1. Hodi, F.S., et al., Improved survival with ipilimumab in patients with metastatic melanoma. N Engl J Med, 2010. 363(8): p. 711–23.

2. Topalian, S.L., et al., Safety, activity, and immune correlates of anti-PD-1 antibody in cancer. N Engl J Med, 2012. 366(26): p. 2443–54.

3. Wolchok, J.D., et al., Nivolumab plus ipilimumab in advanced melanoma. N Engl J Med, 2013. 369(2): p. 122–33.

4. Miceli, R., et al., Sex differences in burden of adverse events in patients receiving immunotherapy. Journal of Clinical Oncology, 2023. 41(16_suppl): p. 2646–2646.

5. Conway, J.R., et al., Genomics of response to immune checkpoint therapies for cancer: implications for precision medicine. Genome Medicine, 2018. 10(1): p. 93.

6. Gaw, N., S. Yousefi, and M.R. Gahrooei, Multimodal data fusion for systems improvement: A review. IISE Transactions, 2022. 54(11): p. 1098–1116.

7. Argelaguet, R., et al., Multi-Omics Factor Analysis—a framework for unsupervised integration of multi-omics data sets. Molecular Systems Biology, 2018. 14(6): p. e8124.

8. Argelaguet, R., et al., MOFA+: a statistical framework for comprehensive integration of multi-modal single-cell data. Genome Biology, 2020. 21(1): p. 111.

9. Furney, S.J., et al., Genome sequencing of mucosal melanomas reveals that they are driven by distinct mechanisms from cutaneous melanoma. The Journal of Pathology, 2013. 230(3): p. 261–269.

10. Ma, Y., et al., Mucosal Melanoma: Pathological Evolution, Pathway Dependency and Targeted Therapy. Frontiers in Oncology, 2021. 11.

11. Mihajlovic, M., S. Vlajkovic, P. Jovanovic, and V. Stefanovic, Primary mucosal melanomas: a comprehensive review. International journal of clinical and experimental pathology, 2012. 5(8): p. 739–753.

12. Newell, F., et al., Whole-genome landscape of mucosal melanoma reveals diverse drivers and therapeutic targets. Nature Communications, 2019. 10(1): p. 3163.

13. Bastiaan, W.H., et al., Intestinal transkingdom analysis on the impact of antibiotic perturbation in health and critical illness. bioRxiv, 2020: p. 2020.06.25.171553.

14. Aitchison, J., The Statistical Analysis of Compositional Data. Journal of the Royal Statistical Society: Series B (Methodological), 1982. 44(2): p. 139–160.

15. Gloor, G.B., J.M. Macklaim, V. Pawlowsky-Glahn, and J.J. Egozcue, Microbiome Datasets Are Compositional: And This Is Not Optional. Front Microbiol, 2017. 8: p. 2224.

16. Liberzon, A., et al., Molecular signatures database (MSigDB) 3.0. Bioinformatics, 2011. 27(12): p. 1739–1740.

17. Subramanian, A., et al., Gene set enrichment analysis: A knowledge-based approach for interpreting genome-wide expression profiles. Proceedings of the National Academy of Sciences, 2005. 102(43): p. 15545–15550.

18. Cantini, L., et al., Benchmarking joint multi-omics dimensionality reduction approaches for the study of cancer. Nature Communications, 2021. 12(1): p. 124.

19. Tenenhaus, M., A. Tenenhaus, and P.J.F. Groenen, Regularized Generalized Canonical Correlation Analysis: A Framework for Sequential Multiblock Component Methods. Psychometrika, 2017. 82(3): p. 737–777.

20. Bady, P., S. Dolédec, B. Dumont, and J.-F. Fruget, Multiple co-inertia analysis: a tool for assessing synchrony in the temporal variability of aquatic communities. Comptes Rendus Biologies, 2004. 327(1): p. 29–36.

21. Hayward, N.K., et al., Whole-genome landscapes of major melanoma subtypes. Nature, 2017. 545(7653): p. 175–180.

22. Chung, Y.J., et al., Iron-Deficiency Anemia Results in Transcriptional and Metabolic Remodeling in the Heart Toward a Glycolytic Phenotype. Front Cardiovasc Med, 2020. 7: p. 616920.

23. Watson, A.S., et al., Association of Immune-Related Adverse Events, Hospitalization, and Therapy Resumption With Survival Among Patients With Metastatic Melanoma Receiving Single-Agent or Combination Immunotherapy. JAMA Network Open, 2022. 5(12): p. e2245596–e2245596.

24. Kobayashi, Scott D., N. Malachowa, and Frank R. DeLeo, Neutrophils and Bacterial Immune Evasion. Journal of Innate Immunity, 2018. 10(5-6): p. 432–441.

25. Sadiku, P., et al., Neutrophils Fuel Effective Immune Responses through Gluconeogenesis and Glycogenesis. Cell Metabolism, 2021. 33(2): p. 411–423.e4.

