## Supplemental Figure 1 for "Integrative Multi-Omics Analysis of Melanoma: Uncovering Pathways Associated with Immunotherapy Outcomes"

### Supplementary Figure S1. Tumor-data derived MOFA Factors are not defining for patient demographics or co-morbidities

**A**

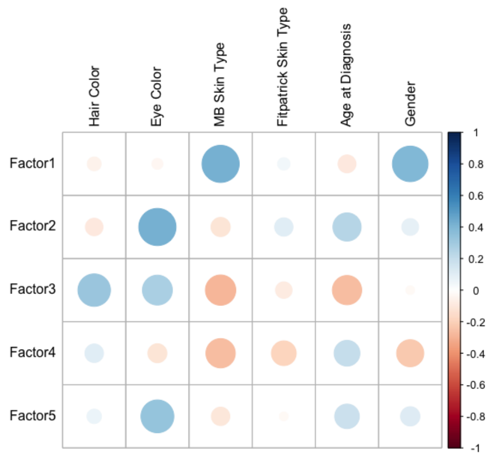


**B**

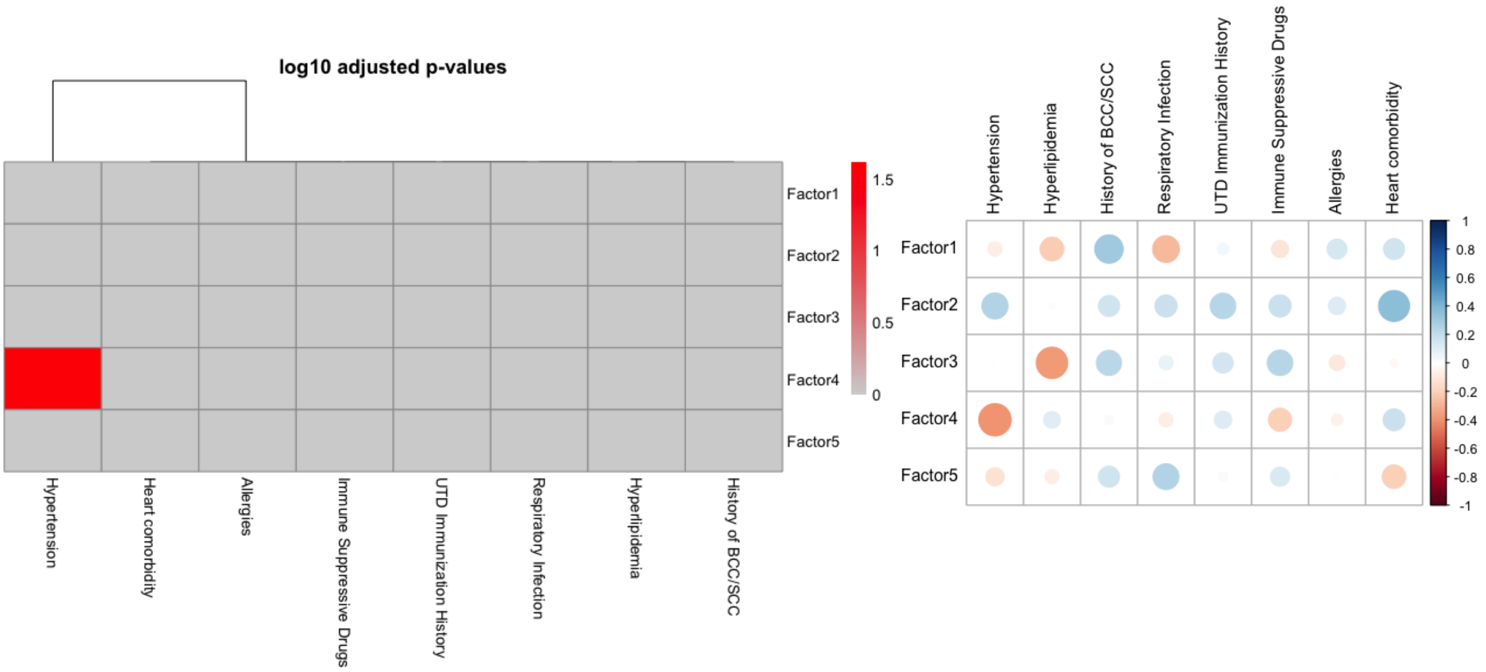
