## Supplementary figures and images for "Integrative Multi-Omics Analysis of Melanoma: Uncovering Pathways Associated with Immunotherapy Outcomes"

### Supplemental Figure 2

#### Supplementary Figure S2. Additional Factor 4 data

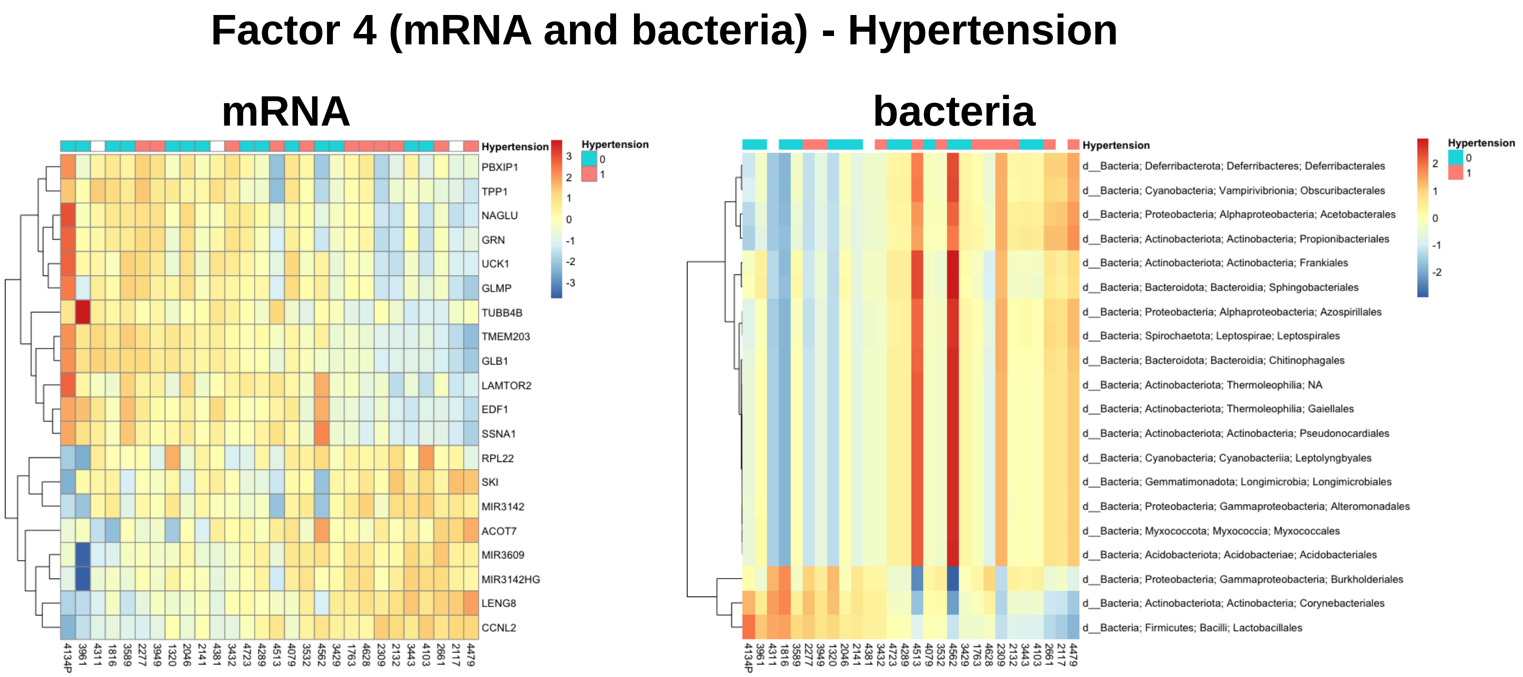
