## Supplemental Figure 3 for "Integrative Multi-Omics Analysis of Melanoma: Uncovering Pathways Associated with Immunotherapy Outcomes"

### Supplementary Figure S3. Factor 5 data

**A**

#### mRNA - Headache

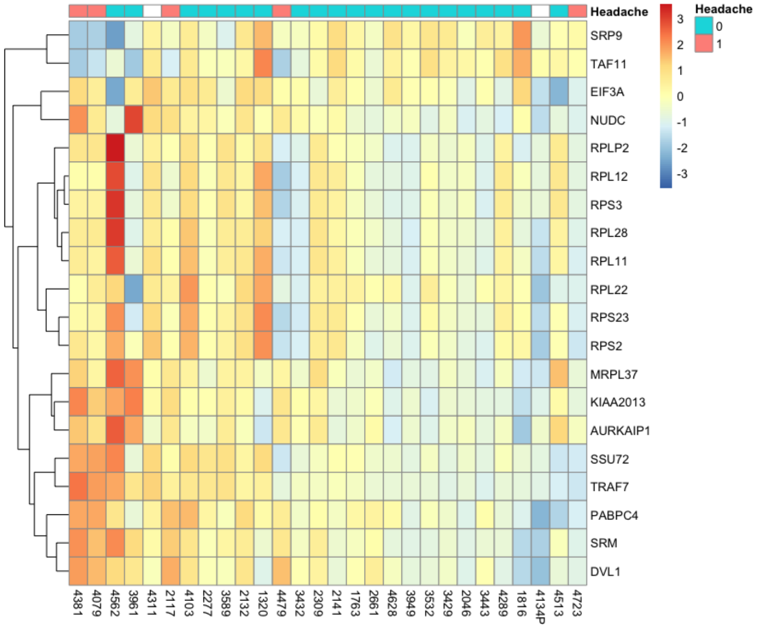


**B**

#### Pathways associated with positive factor weights – Factor 5

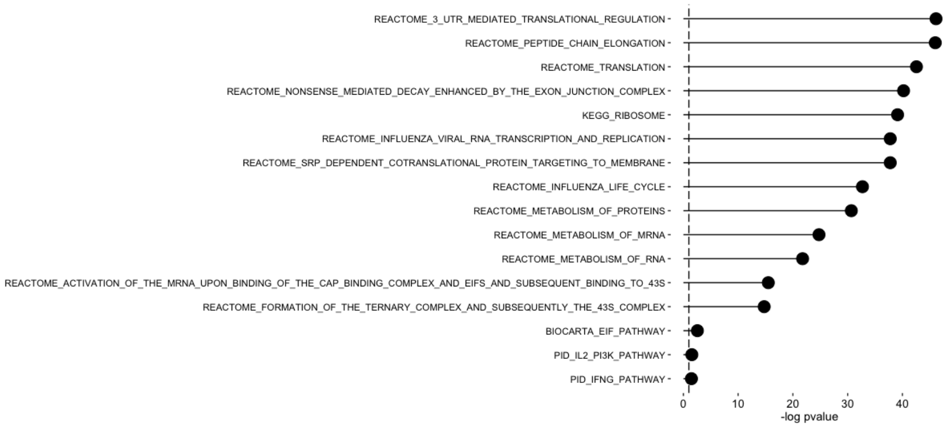


**C**

#### Pathways associated with negative factor weights – Factor 5

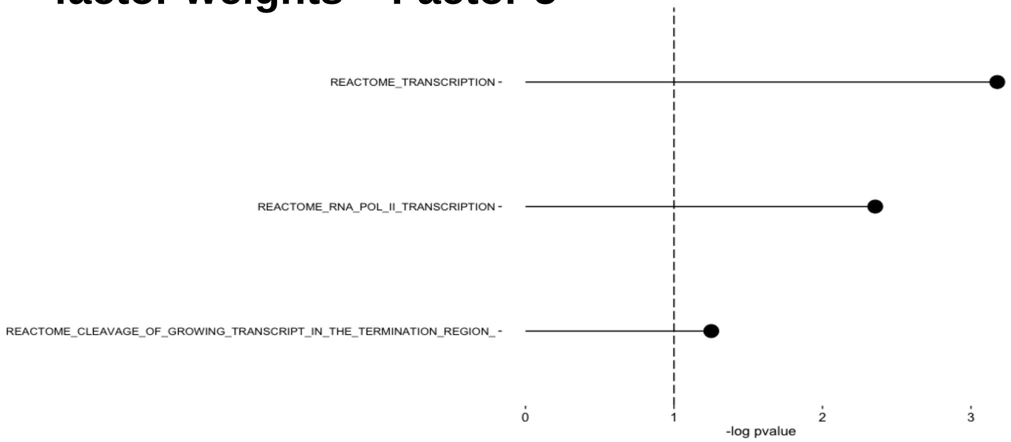
